# Significance of the Wnt canonical pathway in radiotoxicity via oxidative stress of electron beam radiation and its molecular control in mice

**DOI:** 10.1101/2021.03.05.434047

**Authors:** Shashank Kumar, Eram Fathima, Farhath Khanum, Suttur S Malini

**Affiliations:** Molecular Reproductive and Human Genetics Laboratory, Department of Zoology, University of Mysore, Manasagangothri, Mysuru- 570006, Karnataka, India; Defense Food Research Laboratory, Defense Research Development Organisation, Siddharthanagar, Mysore

**Keywords:** Electron Beam Radiation, Wnt canonical pathway, Apoptosis, Oxidative Stress, DNA damage, Radio-toxicity

## Abstract

Radiation triggers the cell death events through signaling proteins, but the combined mechanism of events is unexplored. We intend to investigate how the combined cascade works throughout the radiation process and its significance over radiotoxicity. Thirty adult mice were irradiated with electron beam radiation, and five were served as a control (Non treated). Mice were sacrificed after post 24 hours and 30 days of irradiation. We assessed the oxidative stress parameter and mRNA profile (Liver, Kidney, spleen, and germ cells), sperm viability, and motility. The mRNA profile study established the network of combined cascade work during radiation to combat oxidative response and cell survival. The quantitative examination of mRNA uncovers unique critical changes in all mRNA levels in all the cases, particularly in germ cells. Recuperation was likewise seen in post 30 days radiation in the liver, spleen, and kidney followed by oxidative stress parameters, however not in germ cells. It proposes reproductive physiology is exceptionally sensitive towards radiation, even at the molecular level. It also suggests the suppression of Lef1/axin2 could be the main reason for permanent failure in the sperm function process. Post irradiation likewise influences the morphology of sperms. The decrease in mRNA level of Lef1, Axin2, survivin, Ku70 suggests radiation inhibits the Wnt canonical pathway and failure in DNA repair mechanism in a coupled manner. Likewise, the increase in BAX and BCL2 suggests apoptosis activation followed by the decreased expression of enzymatic antioxidants. In summary, controlled several interlinked cascades execute when body exposure to radiation may further be used in a study to counterpart and better comprehend medication focus on radiation treatment.

## Introduction

Electron beam radiation is a type of ionizing radiation (IR), which can ionize the particle to which it strikes. It is often used in cancer therapy to eradicate the malignant cells, and radiation interactions to molecular proteins are always hostile. Radiation has been proven to be a known factor to manipulate the physiology of the cells by transmitting its energy and altering the cellular metabolism by producing ROS (reactive oxygen species), which leads to the toxicity of the cells (1). The significant alteration in oxidative stress parameters and its relationship in radiation dose is evident in the present study. The recent development in radiation research has helped investigate the close relationship of different radiation dosages affecting the physiological status. Ionizing radiation is used in a broad range to treat cancer cells. Electron beam transfers their energy to the cancer cells, leading to the disruption of cellular metabolism and denaturing cell proliferating enzyme which causes the death of cancer cells. Radiation is also responsible for the high production of ROS, which causes oxidative stress (1). ROS is a collective term of all oxygen molecules and their derivatives, free electrons on its molecule. Due to the instability of oxygen with free electrons, it tends to release the electron and transfer to a native molecule which it comes across. Molecular oxygen that is di-radical is not reactive, but the incomplete reduction of oxygen molecules leads to ROS production (2). ROS is generally considered as superoxide, peroxide, and hydroxyl radical. The classical finding has proven that ROS has been considered the principal source of oxidative stress mediating pathology because they functionally and structurally disrupt the macromolecules such as nucleic acids, protein, and lipids (3). A marginal amount of ROS is always present in normal cells, but when the amount of ROS is generally high, that situation is considered as Oxidative stress, and that has been implicated with Carcinogenesis(3), Neurodegeneration (4), atherosclerosis(5), Diabetes (6), Aging, etc.

Radiation therapy leads to death to even normal cells when irradiated along with cancer cells. Several mechanisms have been previously elucidated in physiology after irradiation. Currently, radiotherapy combined with targeted therapy has become interested in studies on radiation biology in cancer stem cells (CSCs), apoptosis, reactive oxygen species (ROS), and DNA damage repair. To elucidate the signaling pathways involved in radioprotection towards normal cells and radiosensitization towards cancer cells, which is now of primary importance in radiation research (7). Dysregulation of Wnt signaling leads to cancer tumorigenesis, which triggers a series of molecular changes, including oncogenes’ activation and inactivation of suppressor genes. (8). In the present work, we try to evaluate the significant role of Wnt signaling in radiation biology and the mechanism by which Wnt signaling may promote radioprotection.

IR induces apoptosis and necrosis via a series of events, including water radiolysis, which results in generating ROS (11). Unpaired electrons of free oxygen transfer their free radical to DNA molecule, resulting in IR-induced DNA damage (10). Damaged DNA or excessive ROS initiate Bax and bcl2 upregulation, which induces apoptotic signaling pathways leading to cell death (11).

4R’s states the success or failure of standard clinical radiation treatment: i.e., repair of DNA damage, redistribution of the cell cycle, repopulation of tumors, and reoxygenation of hypoxic tumor areas (12, 13). We assume that normal cells protect themselves during IR exposure by reducing ROS-induced damage, including hypoxia, increasing DNA damage tolerance, ROS scavenger, the antioxidant system’s activity, and enhancing DNA repair mainly by activating intracellular pro-survival and anti-apoptotic signaling pathways. Hence, we try to connect these mechanisms of the Wnt signaling pathway and other related radioprotection pathways in detail. IR not only changes the cells but also remodels the microenvironment of the cells. In summary, Wnts and many other co-activators are induced in both cells because of IR exposure, thus inhibit unwanted apoptosis, suggesting the role of Wnt to protect from IR-induced damage. Hence, to improve radiation therapy’s efficacy in cancer cells and save the normal cells during radiation therapy, Wnt signaling can be an appropriate target.

We aim to monitor these changes in various doses with their radiotoxicity and genomic stability to understand how electron beam radiation behaves in different physiological systems. Hence in this regard, we try to evaluate the radiotoxicity and genomic instability in swiss albino mice due to electron beam radiation at three various doses.

## 1 Materials and Methodology

### 1.1 Group Distribution

35 adult Swiss albino mice having an average weight of 25-30gm (Aged 6 to 8 months old) were taken for the study, and seven groups were made in the arrangement of 5 mice per group. The 1st group served as the control, whereas the other six groups were arranged further based on their dose. Three doses are 4gy, 10gy, and 12gy (Per dose two groups) respectively, which were given to 6 groups in which 1st three consecutive groups were kept for 24 hours to check the immediate effect and the other three consecutive groups were kept for the recovery period.

### 2. rradiation

Irradiation of mice was done under Siemens linear Accelerator at HCG Bharath Cancer Hospital, Mysore. Swiss albino mice were kept in the cage made up of polypropylene 4×4×4 cm^3^ cage. Animals were divided into six groups according to the Table. Each group contains five animals each. These animals were irradiated to 4Gy, 10Gy, and 12Gy at 6 Mev radiation dosages.

### 3. Schematic representation of work done

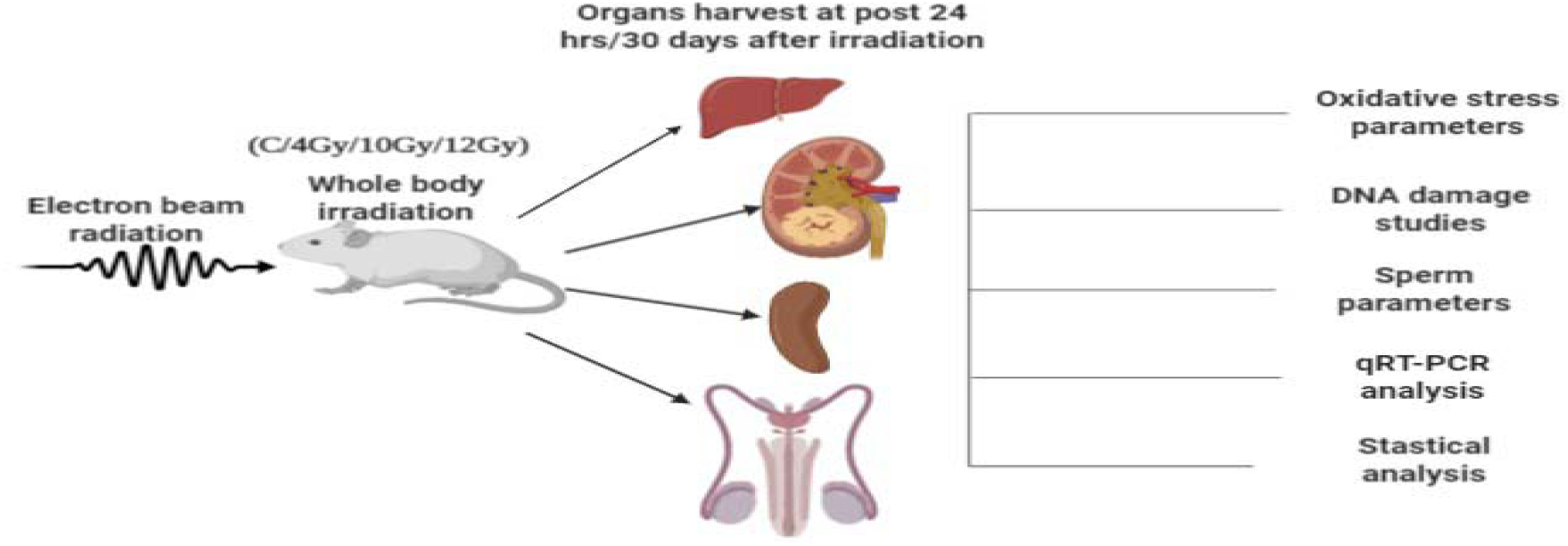

#### 3.1 Preparation of Sample

Mice were sacrificed with the fulfillment of ethical parameters. Spleen, Liver, and Kidney were removed and kept in the PBS with pH-7.4. 10mg of tissue from each group were homogenized with 1ml of PBS and kept centrifugated at 3000 rpm for 10 minutes. The supernatant was used to evaluate biochemical parameters.

#### 3.2 Isolation of Germ cells

Decapsulated testis was minced finely and filtered with a clean muschin cloth. Filtered cells were incubated for 30 minutes in 10ml of RPMI-1640 and collagenase (1mg/ml). After incubation, tubes were centrifuged at 3000rpm for 10 minutes. Discard excess supernatant and keep half of the supernatant to mix. Take the gradient of 2ml each of 37% and 55% percoll was prepared in HBSS. (Modified method of Chang et al., 2011) Cells were mixed in the percoll gradient and kept for 15 minutes. The mixture was centrifuged at 500 x g for 20 minutes for separation of Germ and Somatic cells. Most of the germ cells will be separated in 37% of the percoll gradient (13).

#### 3.3 Preparation of Sample for RNA isolation

Tissues were immediately kept in RNAlater solution, Sigma Aldrich, USA after dissection and were kept -20^0^ storage.

#### 3.4 Evaluation of Oxidative Stress Parameters

##### Superoxide Scavenging Activity

Superoxide dismutase activity has been estimated by the modified method of (14). The 100µl supernatant of homogenized tissue in PBS was taken as the sample. The reaction mixture, Phosphate buffer(0.1 M pH-7.4), α-methionine (20 mM), Hydroxyl Amine Hydrochloride (10 mM), EDTA (50µM), Triton-X (1%), Riboflavin (100 µM) were added to the sample. Without sample serves as a control and without riboflavin serves as blank and the spectrophotometer observed the OD.

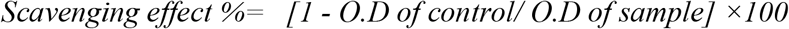

##### Catalase activity

Catalase (CAT) activity was measured by adding 50 µl of the supernatant to a 3 ml reaction mixture containing 8.8 mM H_2_O_2_ (1.5%) and 0.1 M sodium phosphate buffer (pH 7.4), and the absorbance at 240 nm was monitored for 3 min. The decrease in H2O2 concentration was expressed as mmol of H_2_O_2_ decomposed/min/mg of protein. (15)

##### Lipid peroxidation

Lipid peroxidation (LPO) in the homogenate was measured by the thiobarbituric acid reactive substances (TBARS) method (16). 250 µl of the homogenate was mixed with 1.5 ml of each trichloroacetic acid (TCA) (20%) and TBA (0.6%). The mixture was incubated in a boiling water bath for 30 min, cooled, and 2 ml of butanol was added, mixed by vortexing, and centrifuged. The color of the butanol layer was read at 535 nm in a spectrophotometer.

##### Total antioxidant capacity (TAC) in the given sample

The total antioxidant capacity is measured by the modified method of the Phosphomolybdic method. (17)

###### NO scavenging activity

At physiological pH, nitric oxide generated from aqueous Sodium nitroprusside (SNP) solution interacts with oxygen to produce nitrite ions, which may be quantified by the Griess reaction. The activity was measured according to the Griess reaction. One milliliter of 5mM Sodium nitroprusside (SNP) solution was added to 100µl of the homogenized irradiated tissue sample; these reaction mixtures were incubated for 1hr at 27°Cand diluted with 1.2ml of Griess reagent (1% sulfanilamide in 5% H3PO4 and 0.1% naphthyl ethylene diamine dihydrochloride). The absorbance of the chromophore was read immediately at 550nm. (18).

###### Protein Estimation

Total protein was measured according to the (19) method keeping BSA as standard.

###### Sperm motility and vitality assay

Epididymis was surgically removed and cut carefully in 0.9% of NaCl solution. Numbers of Sperms were observed and further categorized according to their motility, which was Rapid progressive (RP), Slow Progressive (SP), Non-progressive (NP), and Immotile (IM). Eosin stain was used to evaluate the vitality of sperm in mice. 0.2% eosin stain was used to stain the sperm and observed immediately after 10 minutes of incubation at Room Temperature (RT). Viability has been shown in a Viable and non-viable stage.

###### DNA Damage

DNA was extracted from Wizard Genomic DNA Purification Kit, Promega, and the USA from liver and kidney tissues according to the given protocol. Isolated DNA was checked for purification at 260/280 nm ratio by spectrophotometer. Gel electrophoresis of isolated DNA was performed on 1.2% Agarose gel and visualized under UV transilluminator.

#### 2.6 RNA isolation and Quantitative RT-PCR analysis

Total RNA was extracted by Trizol method (20), and 2000ng of Total RNA was converted to cDNA by Prime Script RT reagent, Takara, Japan. The primer has been designed by Getprime software (https://gecftools.epfl.ch/getprime/). The primers are as mentioned in Table. 2.

Further cDNA was amplified with the above-given primer, and Real-Time PCR was performed on BioRad CFX-96 Real-time PCR machine. Results were analyzed by using the ΔΔCt method. (21)

##### Ethical Declaration

The Institutional Animal Ethical Committee has approved the study, the University of Mysore, bearing the ethical number: UOM/IAEC/10/2016. All methods were performed under relative guidelines and regulations.

##### Statistical Analysis

Results are presented as the mean ± standard error (SE) and deviation (SD) of measurements made on five mice in each group (3 in case of mRNA quantification) per experiment and are representative of separate experiments. One-way analysis of variance (ANOVA) followed by *post hoc* with Dunnett T2was used to compare all groups and doses at all times when responses were measured at 24 hours and 30 days. Statistical differences were considered significant when *P* ≤ 0.05 using the SPSS version 22 software (Chicago) and GraphPad Prism 9.0.

## 1. Results

### Radiation damage does affect Wnt canonical pathway in Balb/c Swiss albino male mice

There was a significant decrease in the mRNA level of Lef1 and Axin2, suggesting the effect on the Wnt canonical pathway at post 24 hours and 30 days of irradiation. The relative gene expression of these genes was decreased significantly by increasing the dosage of radiation at 4Gy, 10Gy, and 12 Gy in the liver, kidney, spleen, and germ cells of the testis. At post 24 hours of irradiation, there was a significant reduction in fold change of Lef1 and Axin2 mRNA compared to the control group in all four organs. However, at post 30 days, the irradiated group showed significant recovery in the expression of mRNA levels of Lef1 and Axin2 except for the germ cells (Fig 1: A and B, Table3), suggesting the crucial role of the wnt pathway in the failure of the reproductive profile after radiation exposure.

**Fig 1:**
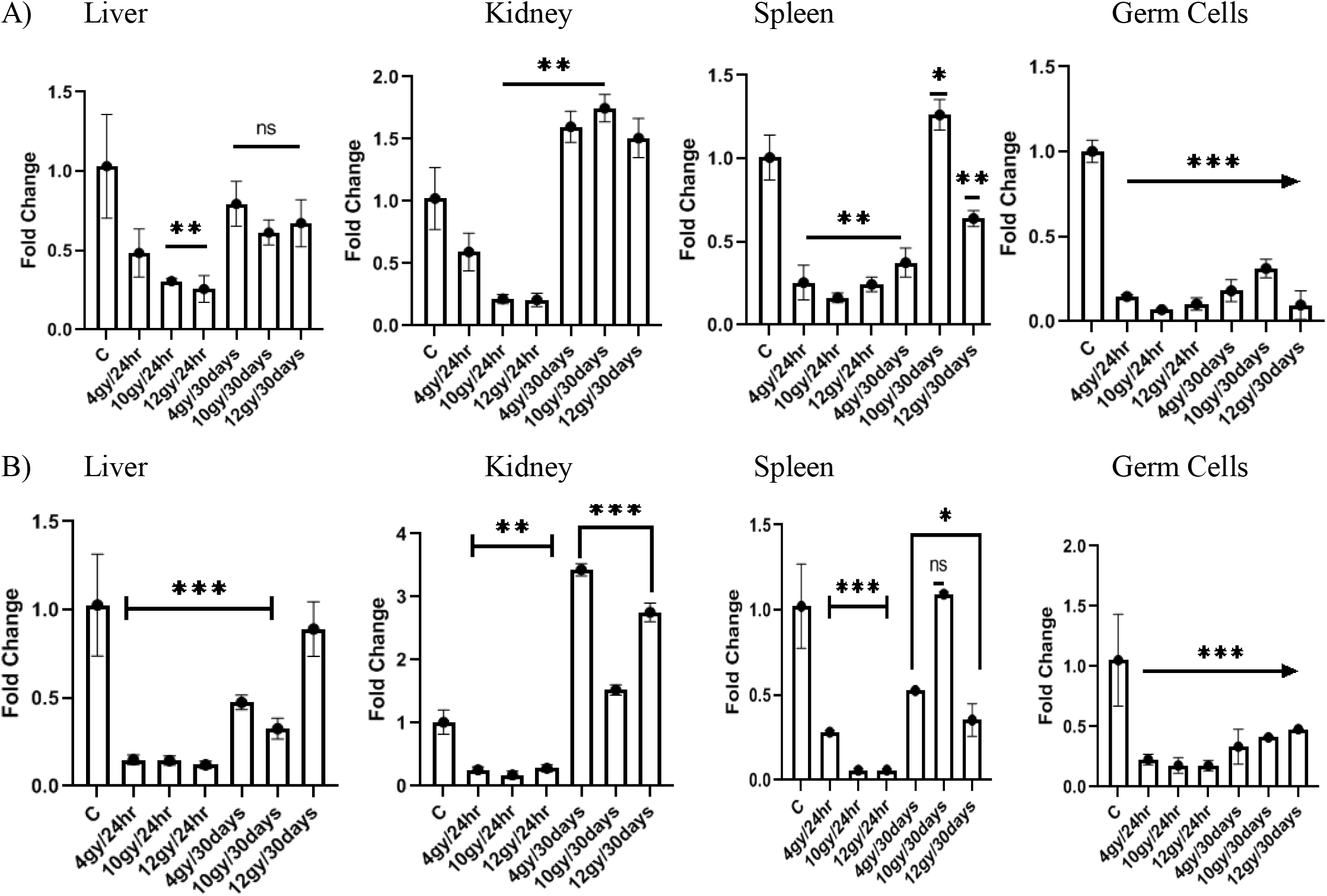
The mRNA level of Lef1 and Axin2, candidates of Wnt-Canonical pathway were measured by RT-qPCR a normalized to GAPDH mRNA in Liver, Kidney, Spleen and Testis (Germ cells) at post radiation of 24hours a 30days. (A) Quantification showed significant Lef1 mRNA reduction in all four tissues (n=3 from 4Gy, 10Gy and 12G mean+ SE, one way Annova, * p<0.05, **p<0.005, ***p<0.001) (B) Quantification showed significant Axin2 mRNA reduction in all four tissues (n=3 from 4Gy, 10Gy and 12G mean+ SE, one way Annova, * p<0.05, **p<0.005, ***p<0.001)

Wnt canonical pathway plays a significant role in spermatogenesis. Since germ cells show poor recovery in Lef1 and Axin2 mRNA level over the post-irradiation period of 30 days suggest its prominent role in permanent failure of reproductive status due to radiation

### Post radiation shows molecular control over apoptosis by regulation of survivin

Bax and Bcl2 are candidate molecules for regulating the intrinsic apoptotic pathway; the up-regulation of Bax and B-cell lymphoma 2 (Bcl2) was evident in our study. The measured level of mRNA by RTqPCR was found to increase in Bax and Bcl2 at post 24hours of irradiation (fig: 2A and B, Table 3), implicating apoptosis was visualized on agarose gel electrophoresis (Fig: 3B). Further, survivin expressed via Wnt canonical pathway inhibits the caspase 3 (22) activity resulting in deregulation of apoptosis and regulating cell survival. The mRNA level of survivin was drastically reduced in post 24hours of post-irradiation, the fold change was found to be approximately negligible in all four tissues (Fig: 2C and Table 3).

In contrast to 24 hours of post-irradiation, 30 days of post-irradiation showed significant recovery of Bax, Bcl2, and Survivin mRNA levels to the average level. The mRNA level was decreased in Bax and Bcl2 compared to post 24 hours of irradiation (Fig: 2A and B, Table 1). As a result of recovery, survivin’s expression was improved significantly to the normal level in the liver, kidney, and spleen. However, in the liver at post 30 days of irradiation, the Bax and Bcl2 were slightly increased as compared to kidney and spleen mRNA levels; on the other hand, poor recovery of Bax, Bcl2, and survivin mRNA was observed in germ cells (Fig: 2A, B, C and Table 3).

### Deregulation in the Wnt canonical pathway enhances the apoptotic event coupled with failure in DNA repair mechanism

Ku70 and Ku80 are two proteins responsible for the non-homologous end-joining pathway (NHEJ), which joins DNA double-strand break. In our study, there was a significant decrease in Ku70 mRNA level in Liver, Kidney, Spleen, and germ cells at post-radiation of 24 hours (Fig: 3A, Table3). The mRNA level of Ku70 was improved during the post-radiation period of 30 days except in germ cells, suggesting the poor recovery in the DNA repair mechanism. Further, the DNA damage was investigated in the irradiated renal and hepatic cell by DNA ladder assay (3b),

Hepatic cells showed severe DNA damage at all dosages over post 24hour of irradiation, but in post 30 days of irradiation, DNA damage was observed only at 4Gy. However, renal cells showed DNA damage only at 4Gy post 24 hours of irradiation (Fig: 3B). Overall, Agarose gel electrophoresis of the Liver and Kidney has shown smear of DNA shear suggest Apoptosis and Necrosis while 4Gy shows long-term radiation effects on the liver but not in the kidney.

### Molecular control of radiotoxicity is coupled with the Wnt canonical pathway via oxidative stress

There was a significant decrease in the mRNA level of SOD1 and SOD2, indicating the effect on superoxide dismutase enzyme at post 24 hours and 30 days of irradiation. The relative gene expression of these genes decreased by increasing the dosage of radiation at 4Gy, 10Gy, and 12Gy, except in the liver, mRNA level was increased at 4Gy after 24 hours post-irradiation. Similarly, the SOD scavenging activity increased at 4Gy and decreased at 10Gy and 12Gy of post-24-hour irradiation (Fig: 4A, 4B and 4E Table 3). The mRNA level of SOD1 and SOD2 were improved during the post-radiation period of 30 days except in germ cells.

Further, there was a significant decrease in Catalase mRNA level in Liver, Kidney, Spleen, and germ cells at post-radiation 24 hours (Fig: 4C, Table1). Likely, Catalase activity was significantly decreased in liver, kidney, and spleen in all dosages of 24 hours and 30 days post-radiation (Fig: 4F Table 3). The catalase level was improved during the post-radiation period of 30 days except in the germ cell.

There was a significant decrease in the mRNA level of iNOS, indicating the effect on inducible nitric oxide synthase at post 24 hours and 30 days of irradiation. The relative gene expression of these genes was decreased by increasing the dosage of radiation at 4Gy, 10Gy and 12 Gy. Similarly, the NOS scavenging activity decreased at 4Gy, 10Gy and 12Gy of post 24 hours irradiation (Fig: 4D and fig: 4G and Table1). The mRNA level of iNOS was improved during the post-radiation period of 30 days except in germ cells (Fig: 4D). Overall, Mice have shown an elevated SOD scavenging activity in 4Gy and decreased in 10 Gy and 12 Gy. Thirty days of post-irradiation shows a slight recovery in SOD scavenging activity but higher as compared to control. Catalase shows the elevated level in a dose-dependent manner in the liver and kidney.

Whereas, spleen shows downgrade activity in a dose-dependent manner. Minor slight improvement in catalase activity during 30 days of the recovery period was observed. Significant elevation in Nitric-oxide scavenging activity has been seen in 4 Gy irradiated mice in the Liver and Spleen. Thirty days of recovery shows a non-significant slight improvement in NO scavenging activity in Spleen and Liver. Higher dose shows a decline in NO scavenging activity, whereas slight improvement has been shown after 30 days of the recovery period. The elevated LPO level (Fig: 4H) has been observed in all conditions even after 30 days of the recovery period, which was in a dose-dependent manner. Spleen shows a high amount of Lipid peroxidation in a dose-dependent manner. Total antioxidant capacity has been compromised after irradiation in a dose-dependent manner. However, slight recovery has been observed in 30 days of post-radiation (Fig:4I)

### Vulnerability in reproductive profile shows the effect of radiation damage even after post-irradiation of 30 days

Germ cells have shown a significant decrease in SOD1, SOD2, iNOS, Cat, Survivin, lef1 and Axin2 (Fig 1A,1B, Fig 2A, 2B, 2C, Fig 3A, 4A, 4B, 4C and 4D). Fig 5A and B shows failure in Sperm viability and motility even after post-radiation. Fig 5C shows different changes in sperm morphology. It suggests deficient reproductive physiology. The decrease in lef1, axin2 and survivin suggest a failure in the Wnt canonical pathway may be the primary reason for the poor condition of germ cell.

**Fig 2:**
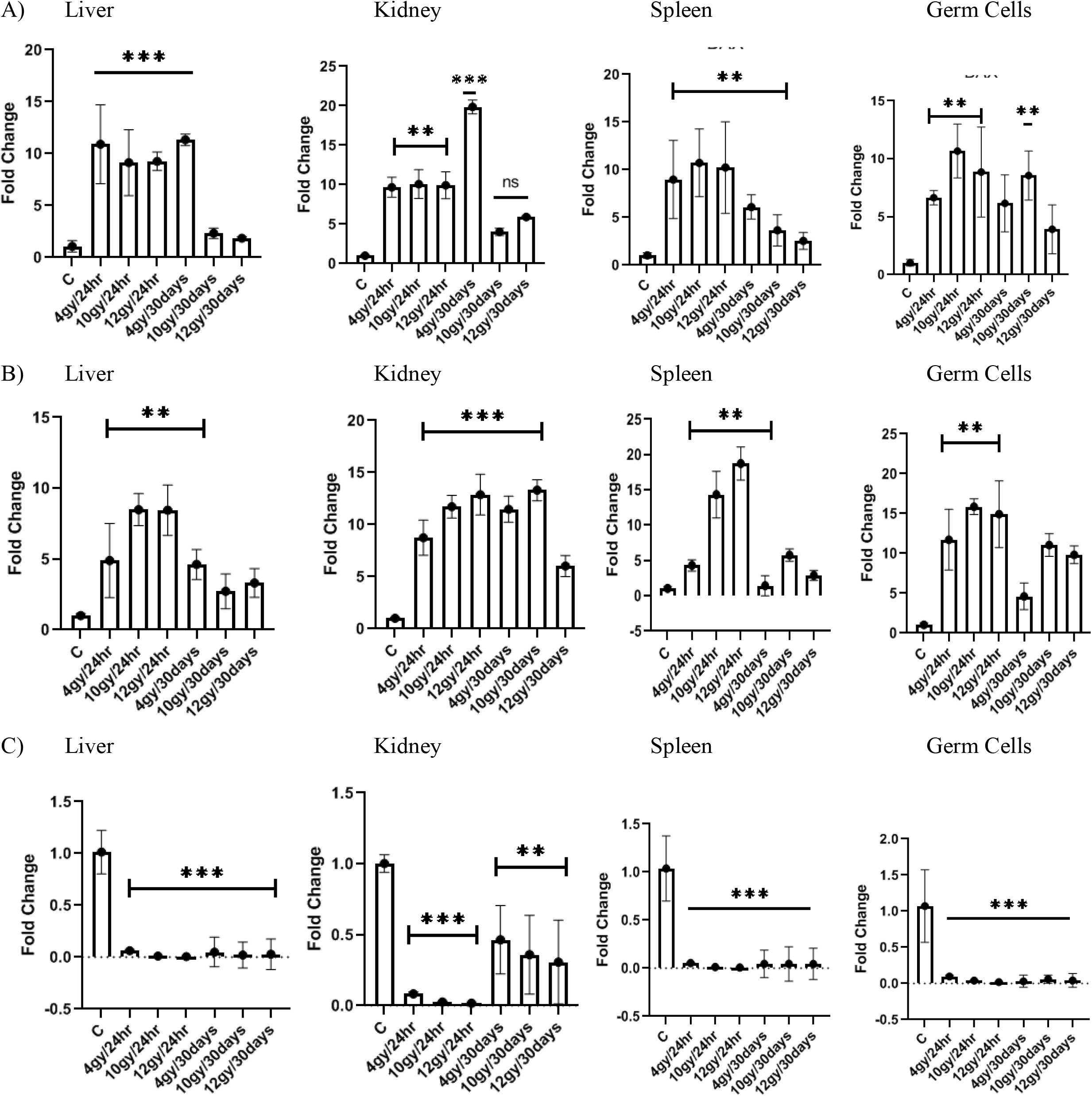
The mRNA level of Bax, Bcl2 and Survivin, were measured by RT-qPCR and normalized to GAPDH mRNA Liver, Kidney, Spleen and Testis (Germ cells) at post radiation of 24hours and 30days. A) Quantification showed significant Bax mRNA reduction in all four tissues (n=3 from 4Gy, 10Gy and 12Gy; mean+ S one way Annova, * p<0.05, **p<0.005, ***p<0.001) B) Quantification showed significant Bcl2 mRNA reduction in all four tissues (n=3 from 4Gy, 10Gy and 12Gy; mean+ S one way Annova, * p<0.05, **p<0.005, ***p<0.001) C) Quantification showed significant Survivin mRNA reduction in all four tissues (n=3 from 4Gy, 10Gy and 12Gy; mea SE, one way Annova, * p<0.05, **p<0.005, ***p<0.001)

**Fig: 3.**
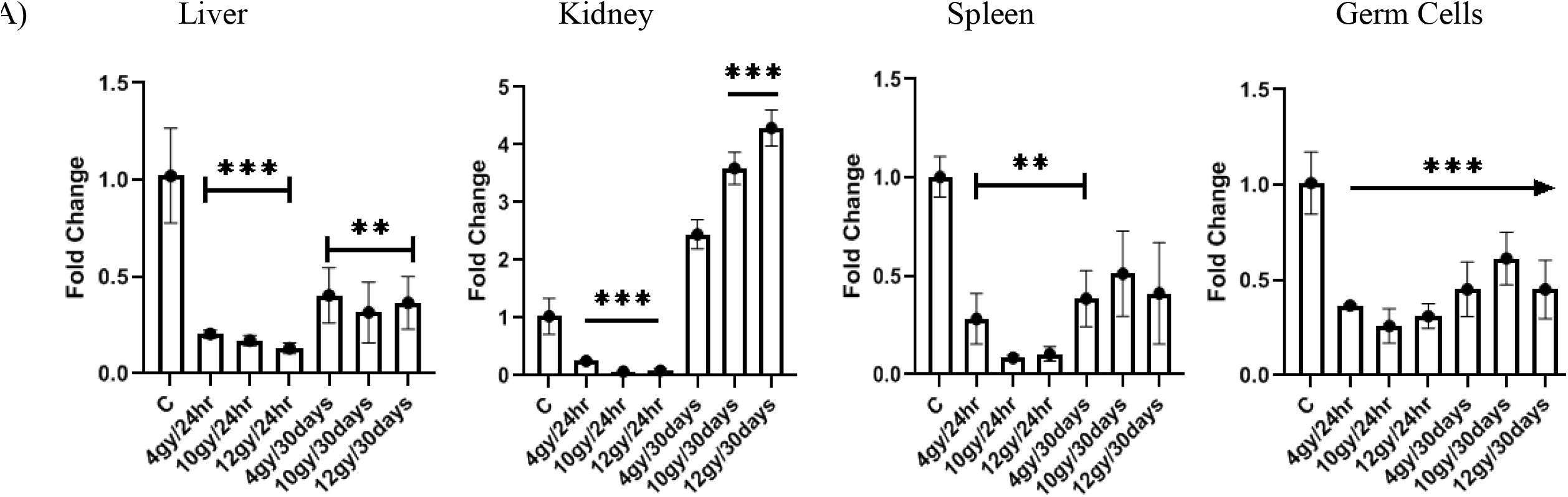
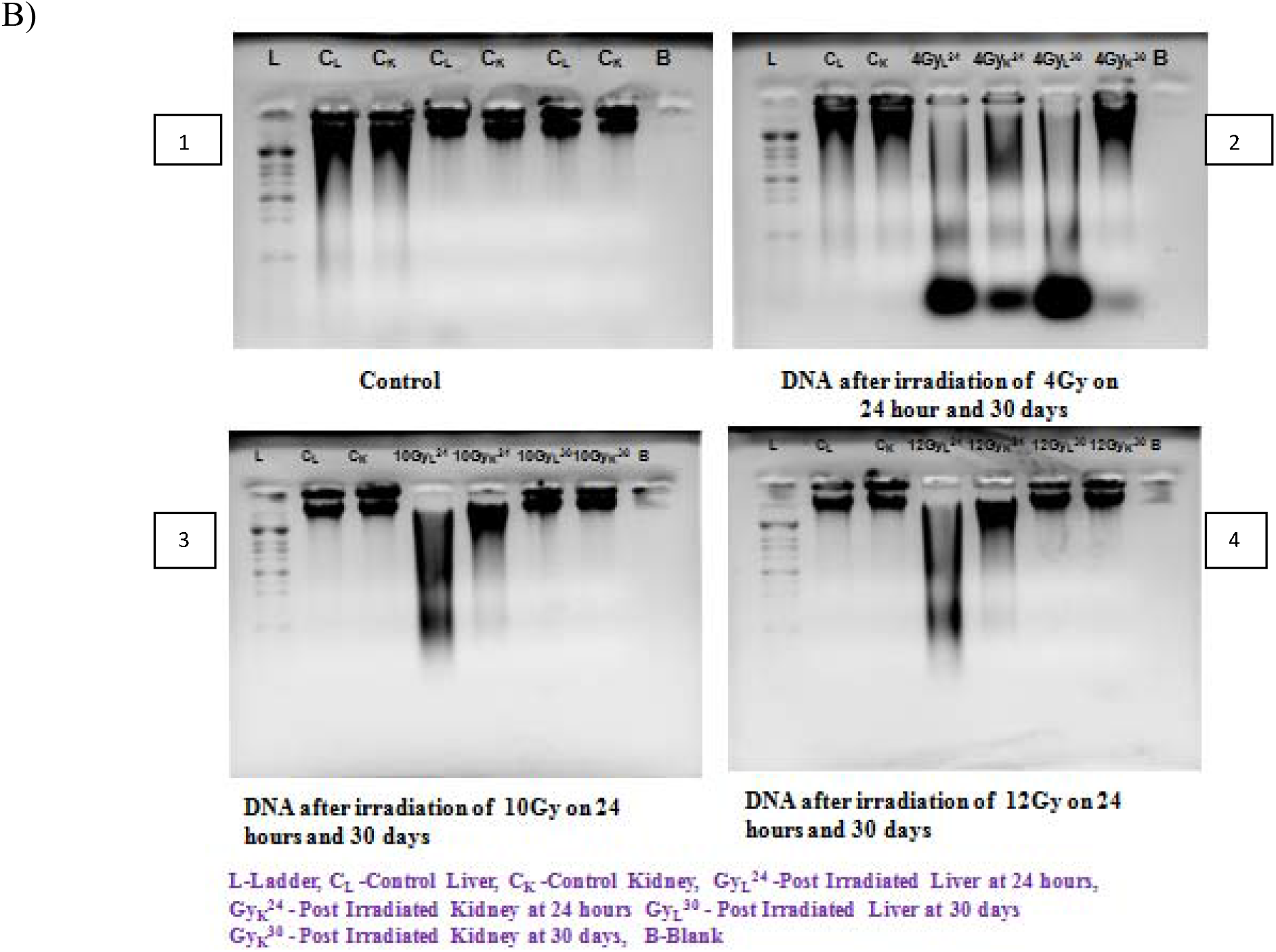
The mRNA level of Ku70 was measured by RT-qPCR and normalized to GAPDH mRNA in L ver, Kidne Spleen and Testis (Germ cells) at post radiation of 24hours and 30days, agarose gel electrophoresis image of hepa and renal cells at 24 hours and 30 days post radiation. A) Quantification showed significant Ku70 mRNA reduction in all four tissues (n=3 from 4Gy, 10Gy and 12Gy; mea SE, one way Annova, * p<0.05, **p<0.005, ***p<0.001) B) 1: Agarose gel electrophoresis of Control liver and kidney in triplicates 2: Agarose gel electrophoresis of liver and kidney in irradiated with 4Gy 3: Agarose gel electrophoresis of liver and kidney in irradiated with 10Gy 4: Agarose gel electrophoresis of liver and kidney in irradiated with 12Gy

**Fig: 4.**
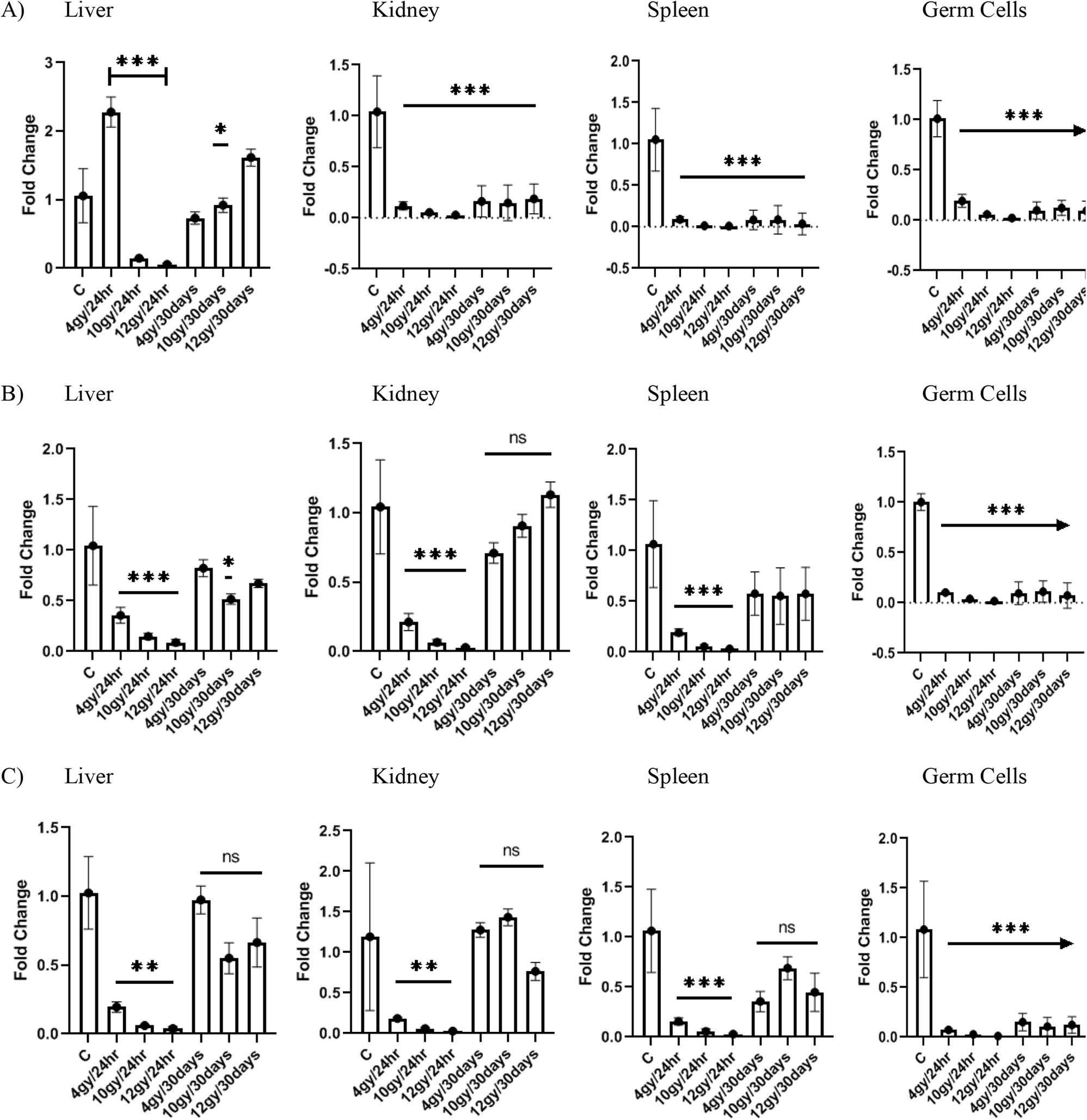
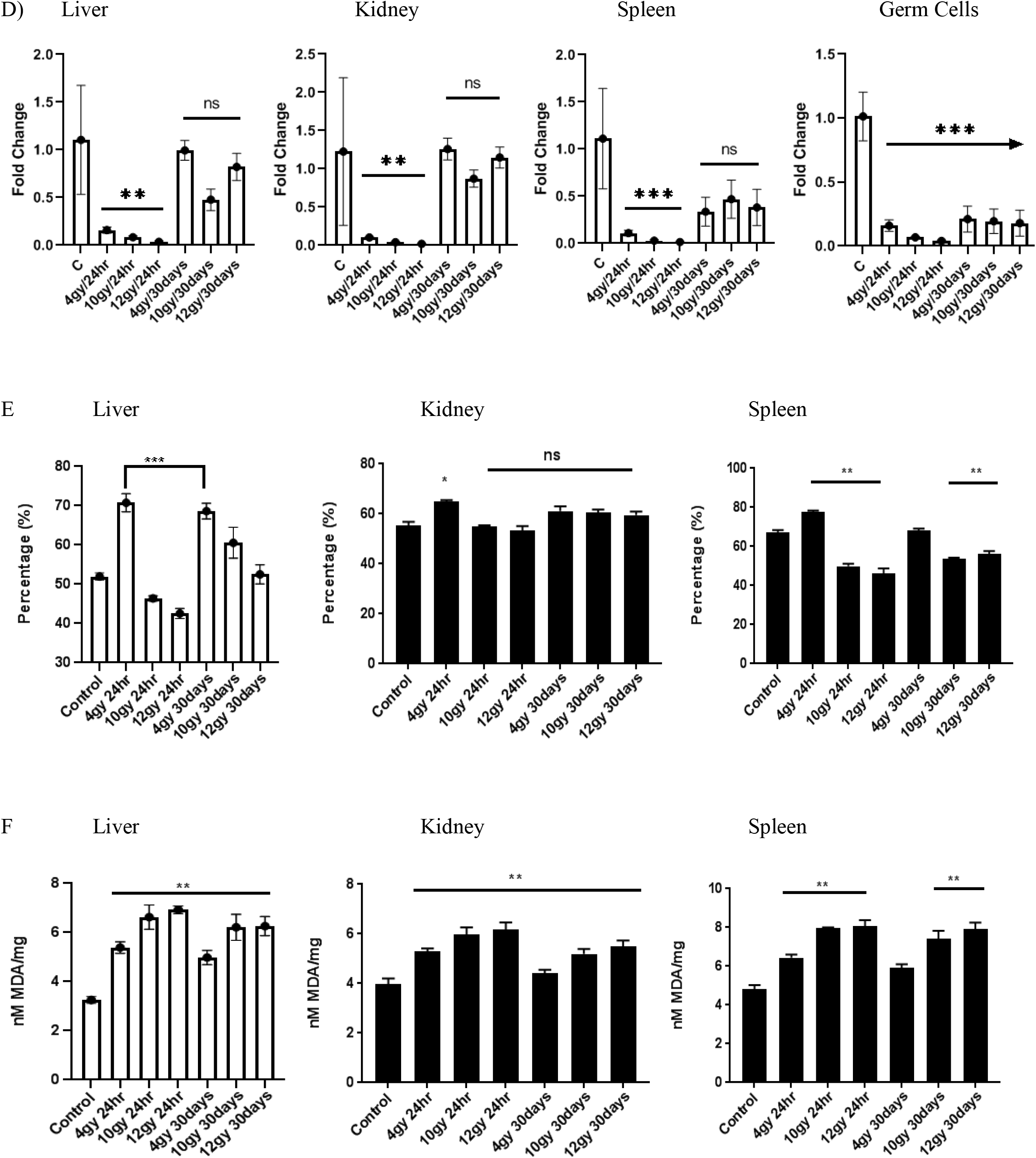
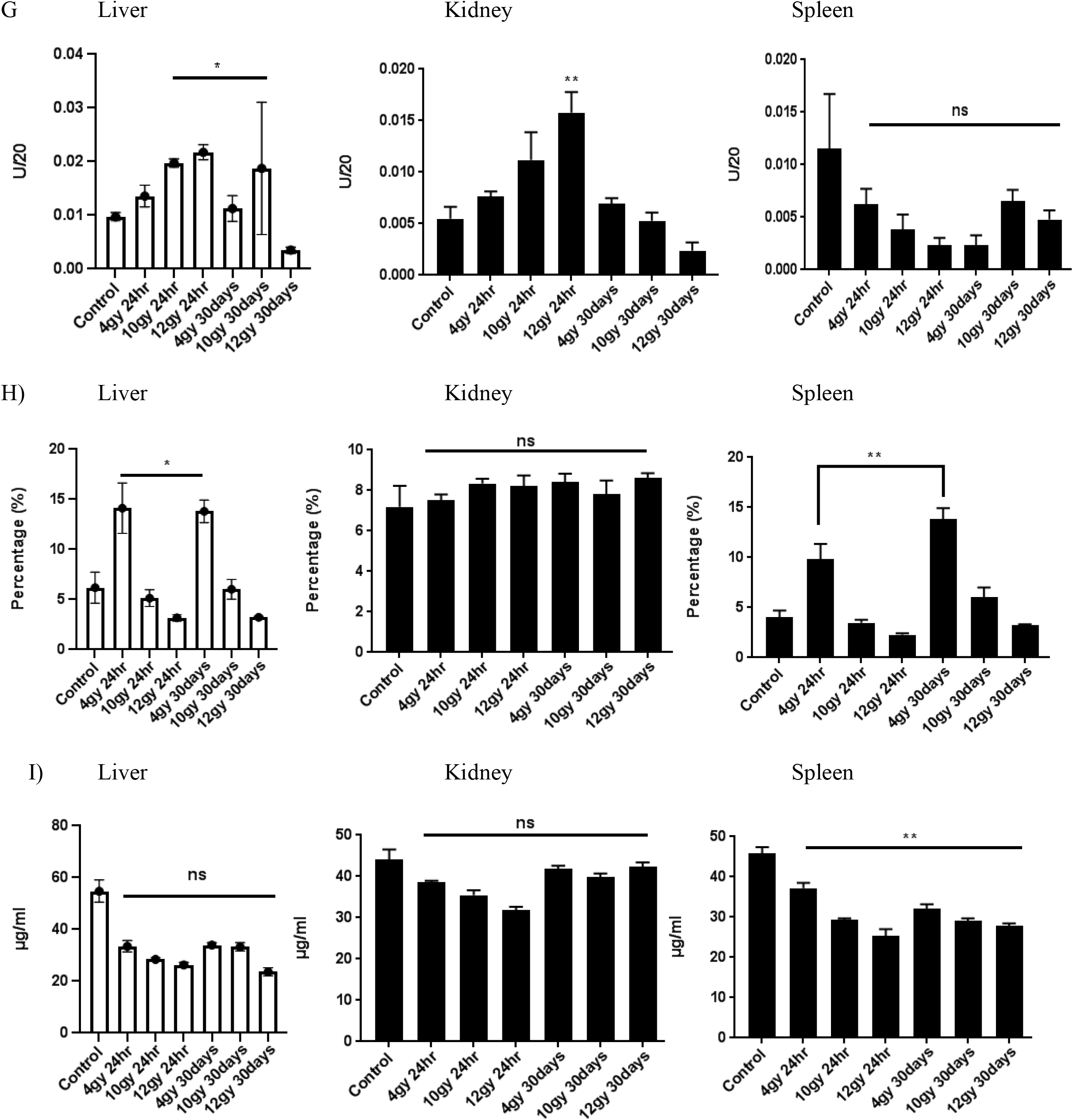
The mRNA level of SOD1, SOD2, Cat and iNOS was measured by RT-qPCR and normalized to GAPDH mRN in Liver, Kidney, Spleen and Testis (Germ cells) at post radiation of 24hours and 30days, SOD scavenging activi Catalase activity, NO scavenging activity, Lipid peroxidation and TAC level was observed. (A) Quantification showed significant SOD1 mRNA reduction in all four tissues (n=3 from 4Gy, 10Gy and 12G mean+ SE, one way Annova, * p<0.05, **p<0.005, ***p<0.001 (B) Quantification showed significant SOD2 mRNA reduction in all four tissues (n=3 from 4Gy, 10Gy and 12G mean+ SE, one way Annova, * p<0.05, **p<0.005, ***p<0.001) (C) Quantification showed significant Cat mRNA reduction in all four tissues (n=3 from 4Gy, 10Gy and 12Gy; mea SE, one way Annova, * p<0.05, **p<0.005, ***p<0.001) (D) Quantification showed significant iNOS mRNA reduction in all four tissues (n=3 from 4Gy, 10Gy and 12G mean+ SE, one way Annova, * p<0.05, **p<0.005, ***p<0.001) Antioxidant activity reduction in all three tissues (n=5 from 4Gy, 10Gy and 12Gy; mean ± SE, one way annova, p<0.05, **p<0.005, ***p<0.001) (E) SOD scavenging activity, (F) Lipid Peroxidation activity, (G) Catalase activi(H) NO scavenging activity, (I) Total antioxidant capacity

**Fig: 5.**
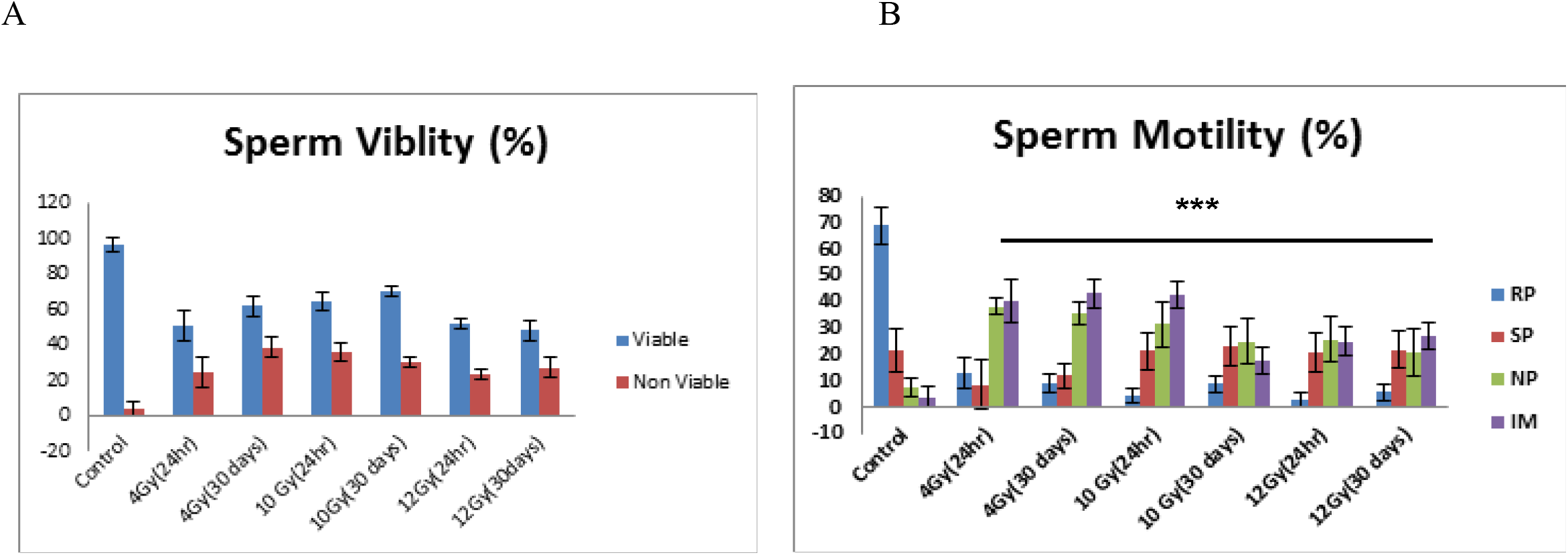
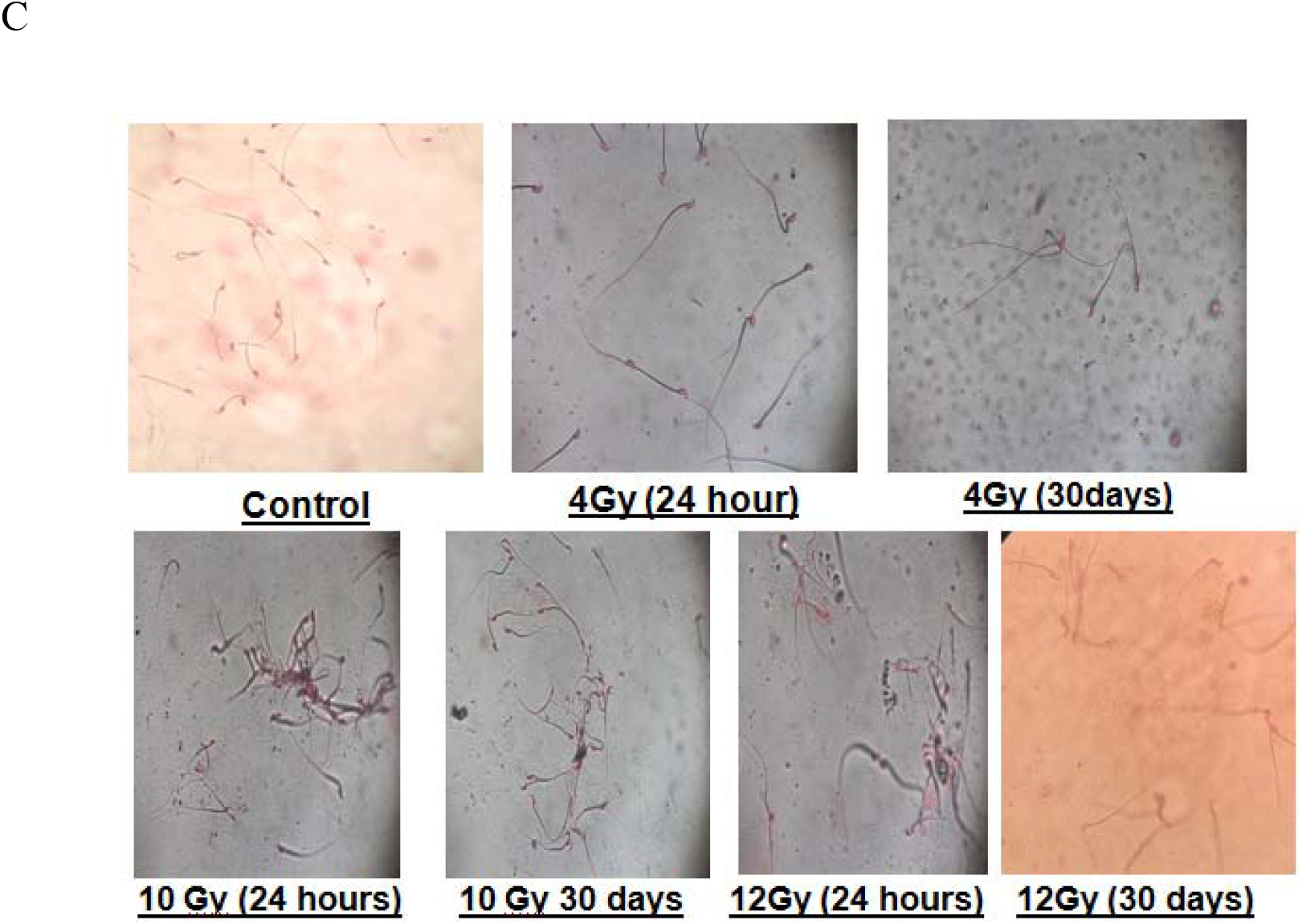
The (A)sperm motility, (B) viability and (C) morphological changes has been observed at post radiation 24hours and 30days, SOD scavenging activity, Catalase activity, NO scavenging activity, Lipid peroxidation and TA level was observed. (n=5 from 4Gy, 10Gy and 12Gy; mean+ standard deviation, one way Annova, * p<0.0 **p<0.005, ***p<0.001)

## Discussion

### Wnt canonical pathway plays a vital role during radiation for cell survival

It is a well-known fact that IR induces ROS and DNA damage, which are the primary cause of cell death. The Wnt canonical pathway in these events has shown its significant role in our study in post-radiation. High levels of ROS irrevocably damage cellular components, including its biomolecules, which further preceded cell death to control the toxic threshold after IR is required to maintain ROS levels (23). We found a similar result, which was reported that in the absence of β-catenin, the expression level of CAT was decreased, and ROS levels were increased, suggesting Wnt canonical pathway plays a significant role in regulating oxidative stress (23). It suggests that β-catenin may protect against oxidative stress damage by controlling the homeostasis between the TCF signaling pathway (mainly proliferative) and the forkhead box O (FOXO) signaling pathway (mainly stress response) (10).

Our study report that there is an underlying coupled and controlled molecular mechanism which executes at radiation exposure via the Wnt canonical pathway by exploring the expression levels of its marker Lef1 and Axin2. The decrease in mRNA level of lef1, axin2 and enzymatic antioxidants leads to inhibition of cell proliferation and cell survival, suggesting the effect of radiation in the Wnt canonical pathway. However, after radiation, Wnt activation can help protect the cells and control the undesirable apoptosis as the recovery has been observed in the expression of its candidate molecules in our study. Along with this, the mRNA of enzymatic antioxidants has been improved at 30 days of post-irradiation. Similarly, (24) proved that the Wnt canonical pathway has a beneficial effect on radiation damage and protecting stem/progenitor cells. Wnt/b-catenin signaling may also suppress apoptosis in the liver, kidney, and spleen by directly inhibiting glycogen synthase kinase 3b (GSK-3b) activity via phosphorylation Wnt coreceptor LPR6 (25). GSK-3b promotes radiation-induced apoptosis by inhibiting pro-survival transcription factors and facilitating pro-apoptotic transcription factors, and GSK-3b inhibitors were shown to protect irradiated small intestine epithelium and hippocampal neurons from apoptosis (26).

### NHEJ pathway shows poor function after irradiation in mice, but recovery has shown after post-radiation of 30days

It is well known that DNA damage gets induced by radiation in 2 ways. These two ways are ROS mediated or direct damage. Ku70/Ku80 is the protein responsible for repairing DNA double-strand breakage in general via the NHEJ pathway (27). Our study suggests that the NHEJ pathway could not repair double-stranded DNA breaks as significantly low Ku70 mRNA level observed as observed in post 24 hours of radiation, resulting in DNA damage; however, recovery has been observed in post 30 days of irradiation. Other studies also showed high doses of radiation often lead to dominant lethal effects, point mutations, and chromosomal abnormalities (28, 29) has shown that abundant low energy of 1 to 20 eV secondary electrons (radical reaction products) play a crucial role in the nascent stages of DNA radiolysis. Ionization of nucleic acid bases has been identified as an initiating step of radiation damage and studied extensively with high energy resolution. (30).

### DNA Damage repair is also coupled with the Wnt canonical pathway

Two major pathways involve DSBs repair: homologous recombination (HR) and non-homologous end-joining (NHEJ). Ku70 inhibits cell proliferation, possibly decreasing B-catenin/Wnt signaling pathway during DNA damage (10). We have observed as Wnt canonical pathway’s candidate molecule and NHEJ candidate molecules are both down-regulated suggest both mechanisms coupled with each other. DNA damage repair mechanism (DDR) trigger cell death when DNA damage is not repaired correctly (31). Previous reports suggested that in the presence of DNA lesions, the DDR is activated, which occurs in three steps: recognition of DNA damage lesions, signal transduction, and DNA damage repair (32). HR is cycle-dependent, which occurs during the G2 and S phases, while NHEJ is cycle-independent (33). x-ray repair cross-complementing 2 (XRCC2), XRCC3, breast cancer type 1 (BRCA1), (BRCA2,) are responsible for HR, whereas KU70 (XRCC6), KU80 (XRCC5), DNA-PKcs, DNA ligase IV (LIG4), and XRCC4 are responsible for NHEJ (43) DSB repair pathway is key to maintaining genome stability. Both HR and NHEJ machinery repair DSBs (34). If damaged DNA cannot be repaired entirely, p53 may trigger apoptotic signaling pathways by upregulating the BH3-only proteins Noxa and Puma (35).

The current study provides experimental evidence that the canonical Wnt-signaling activity mediates normal cells’ sensitivity to DNA damage. Previous studies suggested that Wnt-signaling activity was elevated upon irradiation-induced DNA damage in HSCs and can amplify DNA damage responses in other cell types (36-38). The studies provide a plausible explanation for Wnt signaling maintenance and elimination’s dual effects depending on the level and duration of Wnt-signaling in normal cells. Several studies have proved that the Wnt signaling pathway is essential for self-renewal and regeneration activities upon injury of the ISCs (39-41), supporting the hypothesis that instructed enhancement of Wnt signaling benefits cell regeneration after irradiation. It was shown that Wnt signaling activation resulted in less apoptosis and enhanced proliferation after IR in the gastrointestinal system (42, 43).

### Radiation mediates Apoptotic genes to the programmed cell death via the Wnt canonical pathway

We have seen the up-regulation of Bax and BCL2 mRNA level, which plays a significant role in the intrinsic apoptotic pathway. The intrinsic apoptotic signaling pathway induces various signals (44). An increase in a significant amount of Bax suggests the high number of programmed cell death (45). Bcl-2 family members are essential apoptosis regulators involved in determining cell fate (death or survival). Anti-apoptotic Bcl-2 proteins bind to pro-apoptotic proteins (Bax and Bak), regulating apoptosis (46). BH3-only proteins promote apoptosis by directly activating Bax and Bak, leading to cytochrome c by disrupting the outer mitochondrial membrane (47). Multiple studies suggested a close association between Wnt signaling and apoptotic signaling as the downgrade of the Wnt canonical pathway promotes apoptosis (48). Other studies suggested that FAS-associated factor1 promotes apoptosis by the degradation of β-catenin (49). Our study has observed similar results as upregulation of Bax and Bcl-2 and downregulation of LEF-1, AXIN2 and Survivin, suggesting the close link between Wnt and apoptotic signaling demonstrates that Wnt signaling plays a crucial role in the regulation of radiation mediated apoptosis.

### Oxidative stress, a significant factor in radiotoxicity and inducing/inhibiting combined cascades of NHEJ, Wnt Canonical and Intrinsic apoptotic pathways

Our study found an elevation in lipid peroxidation level and a decrease in TAC level shows severe radiotoxicity in mice. Previous studies have shown a two-fold increase in LPO in liver, kidney, brain, spleen, and testis (50). We also found a 2 to 3 fold increase in LPO in the Liver, Kidney, and Spleen in a dose-dependent manner. LPO leads to progressive loss of membrane integrity, impairment of membrane transport system that disrupts cellular homeostasis (51).TAC status shows the total antioxidant to fight out the ROS in cells. Hence the decrease in TAC shows the weakening in defense mechanism, which leads to cellular toxicity. Superoxide dismutase and catalase are enzymatic antioxidants that breakdown H2O2 to water; it is an essential antioxidant in defense mechanism (48). They work together as superoxide dismutase convert superoxide into peroxide, which catalase further convert peroxide into water. Higher CAT activity seen in irradiated mice indicates enzyme induction due to increased H2O2 formation, a natural adaptive response. The dose-dependent irradiation effect shows a significant increase in oxidative stress during 4 Gy of treatment at 24-hour post-irradiation and a significant decrease in oxidative stress of 10 Gy and 12 Gy treated at 30 days post-irradiation, suggesting the low dosage has a long-term effect compare to high dosage.

Similarly, a dose-dependent decrease in mRNA level of SOD1, SOD2 and Catalase has been observed, suggesting dysregulation in enzymatic antioxidant activity. Nitric oxide scavenging activity reflects the defense mechanism by the cell to protect from nitric oxide free radicals. Nitric oxide free radicals are the byproduct which further created by ROS. It has also abbreviated as RNS, which stands for reactive nitrogen species. It is the first attempt to evaluate the Nitric oxide scavenging activity in irradiated mice, but numerous studies have been done to evaluate Nitric oxide’s function in cell physiology (49). NO is a crucial bioregulatory molecule with several physiological roles, including control of blood pressure, neural signal transduction, platelet function, and antimicrobial and antitumor activities (50). The present study found the significant elevation of NO during a low dose of radiation, which also suggests the adaptive response of a low dosage of radiation, leading to altering the other several biological parameters. Likely, iNOS expression has been severely decreased in a dose-dependent manner suggesting dysregulation of NOS activity leading to various pathophysiology.

### Post radiation studies after 30 days of recovery period shows severe consequences on reproductive physiology

Previous reports have already suggested that radiation leads to permanent failure in sperm motility with a severe decrease in viability and the alteration in morphology. By showing the poor status of Lef1/Axin2 in germ cells after 30 days of post-radiation, we report that Wnt canonical pathway plays a crucial role in spermatogenesis and maintaining its cell biology. Previously, it has been shown that irradiation of the testis with low doses of 1-3 Gy selectively destroys differentiating spermatogonia, resulting in depletion of more advanced spermatogenic cells (50). Stem spermatogonia that are more radioresistant (51) that will not be killed at low doses of radiation maybe because of the Wnt canonical pathway’s up-regulation reported in rats. (49, 50) Although relatively low doses of irradiation caused spermatogenic arrest in certain strains, the surviving cells differentiated and repopulated the seminiferous tubules in other strains with abnormal morphology and decreased motility (52). Radiation leads to an increase in the reactive oxygen species, which are highly toxic to spermatozoa and cause an irreversible arrest to the motility of the sperms, which we also came across with the same observation. The testicular tissues are rich in polyunsaturated fatty acid and are low in antioxidant defense (53). Therefore they are very prone to attack by ROS, which can cause oxidation of proteins, lipids, and DNA leading to cellular damage.

## Acknowledgment and Funding

The author wishes to thank Dr. Vishweshwara and HCG Bharath Cancer hospital staff for providing the irradiation facility. The author wishes sincere thanks to the Board of Research in Nuclear Sciences for providing us non-interrupted funding to carry out this work.

## Summary

Radiation exposure mediates the combined cascade, which is responsible for cell death and cell survival to an extent. Our study has proposed a combined mechanism in which NHEJ and Wnt canonical pathway works together to protect the cells from oxidative stress damage and radiotoxicity. NHEJ pathway repair DNA double breakage during radiation. Furthermore, enhanced apoptotic events increase cell death chances, which inhibit the NHEJ and Wnt canonical pathway. Survivin a candidate molecule express via wnt canonical pathway inhibits caspase3, which further inhibits the apoptotic rate and Wnt pathway positively, correlate with NHEJ pathway. Hence, in conclusion, NHEJ and Wnt canonical pathway suppresses apoptotic event occurs during radiation.

## Author’s contribution

Shashank Kumar conceived and designed the experiment. Shashank Kumar performed Animal handling, harvesting of organs and germ cells, mRNA isolation, cDNA preparation, oxidative stress parameters, DNA ladder, Sperm morphology, viability and motility. Shashank Kumar and Irum Fathima performed RT-qPCR analysis. Shashank Kumar performed the statistical analysis and prepared the manuscript. Suttur S Malini and Farhath Khanum supervised the experiments. Suttur S Malini corrected the manuscript. All authors read and approved the manuscript.

